# Single cell infection with influenza A virus using drop-based microfluidics

**DOI:** 10.1101/2021.09.14.460333

**Authors:** Emma Kate Loveday, Humberto S. Sanchez, Mallory M. Thomas, Connie B. Chang

**Affiliations:** Center for Biofilm Engineering, Department of Chemical and Biological Engineering, Montana State University, Bozeman MT 59717

**Keywords:** Drop-based microfluidics, influenza, single cell, droplets

## Abstract

Influenza A virus (IAV) is an RNA virus with high genetic diversity which necessitates the development of new vaccines targeting emerging mutations each year. As IAV exists in genetically heterogeneous populations, current studies focus on understanding population dynamics at the single cell level. These studies include novel methodology that can be used for probing populations at the single cell level, such as single cell sequencing and microfluidics. Here, we introduce a drop-based microfluidics method to study IAV infection at a single cell level by isolating infected host cells in microscale drops. Single human alveolar basal epithelial (A549), Madin-Darby Canine Kidney cells (MDCK) and MDCK + human *siat7e* gene (Siat7e) cells infected with the pandemic A/California/07/2009 (H1N1) strain were encapsulated within 50 μm radii drops and incubated at 37°C. We demonstrate that drops remain stable over 24 hours, that 75% of cells remain viable, and that IAV virus can propagate within the drops. Drop-based microfluidics therefore enables single cell analysis of viral populations produced from individually infected cells.

## 1. Introduction

Single-cell studies of viral infections enable high-resolution examination of heterogeneous virus populations. Differential selective pressures in both the host cell and virus population lead to variability in virus replication and production that enables antiviral escape, zoonotic spillover events, and changes in virulence or pathogenesis (Dolan, Whitfield, and Andino, 2018). Influenza A virus (IAV) is a negative-strand RNA virus with populations containing high genetic diversity due to its segmented genome, rapid replication rate, and low fidelity RNA-dependent RNA polymerase (RdRp) (Dolan et al., 2018). As such, IAV infection results in a diverse swarm of unique variants exhibiting heterogeneous genotypes and phenotypes (Brooke, 2017; Petrova and Russell, 2018).

Heterogeneous IAV populations can be examined using methods that capture diversity at the single cell level. Such methods include microfluidic techniques to perform single cell transcriptomics (Russell et al., 2019; Russell, Trapnell, and Bloom, 2018; Sun et al., 2020) or isolation of cells using limiting dilutions in well plates (Heldt et al., 2015; Kupke et al., 2020). These studies have identified large variations in total viral mRNA expressed, infectious virus produced, and host transcriptional response to infection. However, isolating single cells using limiting dilutions can limit analysis to only a few hundred cells (Heldt et al., 2015; Kupke et al., 2020; Russell et al., 2019; Russell et al., 2018; Sun et al., 2020).

A promising method to study single cell infection at high-throughput is drop-based microfluidics, which offers the ability to compartmentalize and rapidly assay individual cells (Matuła, Rivello, and Huck, 2020; Mazutis et al., 2013; Prakadan, Shalek, and Weitz, 2017; Xu et al., 2020; Zilionis et al., 2017). A drop-making microfluidic device is used to create microscale aqueous drops that are surrounded by an immiscible oil and stabilized with a biocompatible surfactant (Holtze et al., 2008). Compared to larger scale *in vitro* culturing methods where cells are grown in flasks or well plates, drop-based microfluidics creates millions of discrete bioreactors that house single cells in which viruses can replicate (Fischer et al., 2015; Tao et al., 2015a). This allows for viral replication events to be analyzed independently within these droplet bioreactors. Drop-based microfluidics has recently been applied to study infectivity (Fischer et al., 2015; Tao et al., 2015a) and recombination (Tao et al., 2015b) of the non-enveloped, positive-sense murine norovirus (MNV-1) (Henderson, 2008). Expanding the applicability of drop-based microfluidics to other viruses, such as enveloped viruses that require a broad range of host cells, would greatly increase capacity for single cell virology studies.

In this work, drop-based microfluidics was applied towards culturing and quantifying IAV replication at the single cell level. Three different cell lines, alveolar basal epithelial (A549) cells, Madin-Darby Canine Kidney (MDCK) cells and MDCK + human *siat7e* gene (Siat7e) cells (Chu et al., 2009; Chu et al., 2010), were tested for their viability and ability to support IAV infection in drops compared to bulk tissue culture. Initial one step growth curves in all three cell lines were performed with the A/California/07/2009 H1N1 IAV strain. Both A549 and MDCK cells supported high levels of RNA replication while the Siat7e cells had a stagnant level of viral RNA present over 48 hours (h). Drop stability, cell loading, and viability was tested within 100 µm diameter drops and quantified after 24 h of incubation, a sufficient amount of time for one productive round of IAV infection. The drop radii (*R*) between 0 and 24 h for all three cell lines differed by less than 1.6 µm after incubation demonstrating consistent drop stability. Cell loading, or the number of cells loaded within drops, was quantified and compared using two methods of imaging: a high-speed camera and still images of drops on a hemocytometer. The two methods of measuring cell loading were determined to be incomparable using a mixed effects Poisson model. Quantification of cell loading from the still images was closest to the estimated Poisson loading model for all three cell lines, suggesting its usage as a more reliable method for measuring cell loading compared to the high-speed camera. All cell lines were found to have a minimum mean of at least 75% viability with no significant difference between adherent and non-adherent cell types. The cells were infected with A/California/07/2009 H1N1 at an MOI of 0.1 to compare virus production over time between cells infected on standard tissue culture plates and cells infected and encapsulated as single cells in drops. Viral replication and production was quantified using reverse transcription quantitative polymerase chain reaction (RT-qPCR) and plaque assays. For both A549 and MDCK cells, the increase in detectable viral RNA as measured by RT-qPCR, from 0 to 24 h was similar for both drop and bulk infections. Similar results were observed for measurements of infectious virus produced for both drop and bulk infections of MDCK and A549 as measured by plaque assays. Our findings demonstrate that standard adherent cell lines for propagating and studying IAV can be used in drops, thereby expanding the capabilities to study virus infection at a single cell level, which until recently has only been demonstrated for non-adherent, spinner-adapted host cells (Fischer et al., 2015; Tao et al., 2015a; Tao et al., 2015b).

The drop-based microfluidic methods developed in this work will enable future studies of single cell IAV infections using multiple cell lines. The methods expand the application of drop-based microfluidics from the non-enveloped, positive-sense MNV-1(Fischer et al., 2015; Tao et al., 2015a; Tao et al., 2015b) to include the enveloped and negative-sense IAV, thus broadening the scope of single cell virology assays. Additionally, this work can serve as a blueprint for testing, comparing and implementing drop-based microfluidics methods in future single cell studies. With the recent interest and development of single cell -omic technologies for studying viral infections (Combe et al., 2015; Cristinelli and Ciuffi, 2018; Drayman et al., 2019; Gérard et al., 2020; Guo et al., 2017; Kupke et al., 2020; Liao et al., 2020; Lin et al., 2019; Sen et al., 2012; Timm, Warrick, and Yin, 2017; Zanini et al., 2018), we expect this work to further expand capabilities and increase applications of single cell technologies in virology.

## 2. Material and Methods

### 2.1 Cells and viruses

Human alveolar epithelial cells (A549), Madin-Darby Canine Kidney cells (MDCK) and MDCK + human *siat7e* gene (Siat7e) cells were obtained from ATCC (CCL-185, CCL-34) and Dr. Joseph Shiloach (NIH), respectively (Chu et al., 2009; Chu et al., 2010). A549 cells were propagated in Hams F-12 (Corning) media supplemented with 10% fetal bovine serum (HyClone) and 1X Penicillin/Streptomycin (Fisher Scientific). MDCK and Siat7e cells were propagated in DMEM (Corning) media supplemented with 10% fetal bovine serum (HyClone) and 1X Penicillin/Streptomycin (Fisher Scientific). The IAV virus strain A/California/07/2009 (H1N1) was obtained from Dr. Chris Brooke (UIUC). Stocks of A/California/07/2009 were propagated and titered on MDCK cells.

### 2.2 Microfluidic device fabrication

Microfluidic devices for drop making were fabricated by patterning SU-8 photoresist (Microchem SU-8 50) on silicon wafers (University Wafer, ID# 447, test grade) to create 100 μm tall and 100 μm wide channels. Polydimethylsiloxane (PDMS) (Sylgard 184) was poured onto the wafers at a 10:1 mass ratio of polymer to cross-linking agent according to standard photolithography techniques (Duffy et al., 1998). Air was purged from the uncured PDMS by placing the filled mold in a vacuum chamber for at least 1 h. The PDMS was cured in an oven at 55 °C for 24 h and ports were punched into the PDMS slab with a 0.75 mm ID diameter biopsy punch (EMS Rapid-Core, Electron Microscopy Sciences). The PDMS slab was bonded to a 2-in by 3-in glass slide (VWR micro slides, cat. #48382-179) after plasma treatment (Harrick Plasma PDC-001) for 60 s at high power (30 W) and 700 mTorr oxygen pressure. The drop making devices were made hydrophobic by flowing Aquapel (Pittsburgh Glass Works) through the channels, followed by blowing the channels with air filtered through a GVS ABLUO™ 25 mm 0.2-μm filter (Fisher Scientific) before baking the devices in an oven at 55 °C for 1 h.

### 2.3 Cell encapsulation in drops

Cells were seeded on to T25 flasks or 6-well plates at a density of 1 × 10^5^ cells/cm^2^ and incubated overnight. Cells were prepared for encapsulation by washing 2× with PBS followed by trypsinization to remove cells from the surface of the flasks or plates. The appropriate culture media containing 10% FBS was added to the cells to neutralize trypsinization and cells were collected and centrifuged at 500 × g for 5 minutes. The cell pellet was gently washed with 3-5 mL of PBS and centrifuged a second time at 500 × g for 5 minutes. The cell pellet was gently resuspended in FBS-free media to reach 2 × 10^6^ cells/mL in their designated media. 10 µL of cells were placed on a hemocytometer to visualize and confirm cell concentration and verify the presence of mostly single cells. Cells were loaded into a 3 mL syringe (BD) for injection onto a flow-focusing microfluidic device for encapsulation into 100 µm drops (∼523 pL) (Anna, Bontoux, and Stone, 2003). Drops are stabilized in a fluorinated HFE7500 oil (3 M) continuous phase with a 1.5% w/w solution of PEG-PFPE_2_-based surfactant (RAN Biotechnologies, 008-FluoroSurfactant). The fluids were transferred into the microfluidic device with syringe pumps (New Era NE-1000) controlled by a custom LabVIEW (2015) program at flow rates of 1000 μL/h the aqueous phase and 3000 μL/h for the oil phase. A 1,500 µL total volume consisting of 375 µL of drops containing cells and 1,125 µL of oil, was collected in a 2 mL microcentrifuge tube (Eppendorf). Following encapsulation, drops containing cells were incubated in an open microcentrifuge tube covered with parafilm at 37 °C with 95% relative humidity and 5% CO_2_ (Fischer et al., 2015). Cells were released from drops by removing the majority of the oil phase and adding 1 mL of a 20% w/w 1H,1H,2H,2H-Perfluoro-1-octanol (PFO) in HFE7500 oil to break the emulsion followed by vortexing. Centrifugation at 500 × g for five minutes was used to separate the aqueous cell suspension from the oil phase, to isolate cells for further analysis.

### 2.4 Drop measurements

Drop radii were measured after imaging a drop monolayer on a hemocytometer with a CCD camera (FLIR Grasshopper3) on an inverted microscope (Nikon Ti-U). The drops were added to a hemocytometer at a density where they are arranged in a monolayer with minimal packing. The minimal packing allows the drops to relax in a spherical shape for better image analysis. A custom MATLAB script was used to determine the radii, *R*, at 0 h and 24 h after the cells were incubated. Six images of drops, with a minimum of 300 drops per image per time point, were analyzed. The *R* for all three cell lines before and after incubation. The drop radii data contains values from multiple timepoints post encapsulation, different cell lines, and multiple experimental dates. A linear mixed effects model, with the experimental date as a random factor, was used to analyze the contributions of cell type and timepoint to the data set.

### 2.5 Cell loading and viability

Two methods were used to quantify cell loading in drops. The distribution of cells/drop was determined using still images of a drop monolayer on a hemocytometer as described in 2.4 and footage of the drop formation junction using a high-speed camera (VEO 710L, Phantom). Two high speed videos of the flow focusing junction were recorded at 6,000 frames per second (fps) to capture the formation of individual drops over the course of several frames, with an average of 505 drops counted from each recording. A Poisson model, with cell lines as the mixed effects, the date as the random effect, and the measurement method as a two-way interaction was applied to determine if the cell encapsulation distributions were a function of the cell line or measurement used. An adjusted p-value was then determined using a multiple comparison across the cell lines and measurement methods. Following encapsulation, drops were incubated at 37 °C with 95% relative humidity and 5% CO_2_. Cells were released from drops as previously described. The cells in the aqueous phase were collected and pelleted at 1500 × g for five minutes. The cell pellet was resuspended in 200 µL PBS. Cells were diluted 1:5 in PBS + 10% trypan blue stain for a final volume of 50 µL to determine cell viability. An unpaired t-test was used to determine if cell viability was significantly different between the cell lines.

### 2.6 IAV infection of adherent cells (A549 and MDCK)

Cells were seeded onto a 6-well plate at a concentration of 1 × 10^6^ cells/well and infected with IAV H1N1 at an MOI of 0.1 in infection media. The infection media consisted of Hams F-12 or DMEM supplemented with 1 mM HEPES (HyClone), 1X Penicillin/Streptomycin (Fisher Scientific), 0.1% Bovine Serum Albumin (BSA) (MP Biomedical) and 1 µg/ml of TPCK (tolylsulfonyl phenylalanyl chloromethyl ketone)-trypsin (Worthington Biomedical) (Weingartl et al., 2010). Briefly, cells were washed with 1X Phosphate Buffered Saline (PBS) (Corning) and incubated with 200 µL of virus inoculum for 1 h. The inoculum was removed and replaced with 1.5 mL fresh infection media for another 1 h incubation. Infection media was removed, cells were washed with PBS, and the infected cells were detached from the plate by trypsinization. Infected cells were processed as described in 2.3 and resuspended in encapsulation media containing either Hams F-12 or DMEM and 1 mM HEPES, 1X Penicillin/Streptomycin, and 0.1% BSA (MP Biomedical). The infected cell suspension was split into two populations with one population replated as a bulk control and the other encapsulated in 100 µm drops. Bulk and drop infections were incubated at 37°C with 95% relative humidity and 5% CO_2_, and frozen at - 20 °C at 0 and 24 h post infection (hpi).

### 2.7 IAV infection of suspension Siat7e cells

To infect the Siat7e suspension cells at an MOI of 0.1, 1 × 10^7^ cells were pelleted at 500 × g for 5 min in a 50 mL conical centrifuge tube then resuspended in 200 µL of virus inoculum in DMEM supplemented with 1 mM HEPES (HyClone), 1X Penicillin/Streptomycin (Fisher Scientific), 0.1% Bovine Serum Albumin (BSA) (MP Biomedical) and 1 µg/ml of TPCK-trypsin (Weingartl et al., 2010). The 50 mL tube containing the cells and virus inoculum was shaken at 90 rpm for 1 h. The cells were pelleted at 500 × g for 5 min and the inoculum was removed and replaced with fresh DMEM infection media (10 mL) for bulk infections or DMEM encapsulation media as described in 2.6. Infected cells were either encapsulated in 100 µm drops, or replated as a bulk control. For bulk infections, a 1-mL volume of cells was added to each well of a 6-well plate (2 total) and placed on the shaker. Encapsulated cells were processed as described in 2.3. Bulk and drop infections were incubated at 37°C with 95% relative humidity and 5% CO_2_, and frozen at -20 °C at 0 and 24 hpi.

### 2.8 M gene abundance by RT-qPCR

RNA was extracted from cell suspensions using the QIAGEN QIAmp viral RNA mini kit. RNA copy number of the IAV Matrix protein gene (M gene) was determined using a Taqman RT-qPCR assay (Loveday et al., 2021). Amplification primer sequence was originally described by Shu *et al* (Shu et al., 2011): M gene Forward 5’-GACCRATCCTGTCACCTCTGAC-3’, M gene Reverse 5’-AGGGCATTCTGGACAAATCGTCTA-3’. The sequence of M gene Taqman probe was: 5’-/FAM/ TGCAGTCCTCGCTCACTGGGCACG/BHQ1/-3’. Working stocks of the primers and probe (Eurofins Operon) were prepared at 25 µM and 10 µM, respectively, for use in the RT-qPCR reaction. Samples were amplified using a SuperScript III Platinum One-Step RT-qPCR kit (Invitrogen 11732-020) with a final reaction volume of 25 µl. Each reaction mix contained 400 nM M gene Forward and Reverse primers, 200 nM M gene Taqman probe, 0.05 µM ROX reference dye, 5 U/µl SUPERase RNase Inhibitor (Invitrogen AM2694), and 2.5 µl of RNA template. Thermocycling was performed in a RT-qPCR machine (Quantstudio 3, Applied Biosystems) with the following cycling conditions: 1 cycle for 30 min at 60°C, 1 cycle for 2 min at 95°C, and 40 cycles between 15 sec at 95°C and 1 min at 60°C.

### 2.9 Plaque Assay

Post-infection IAV titers were determined by plaque assay on MDCK cells seeded in 6-well plates at 1 × 10^6^ cells per well. Viral supernatant sampled from infected cells at 0 and 24 hpi was serially diluted in DMEM media with 1 mM HEPES (HyClone), 1X Penicillin/Streptomycin (Fisher Scientific), 0.1% BSA and 2 µg/ml of TPCK-trypsin (Worthington Biomedical). Overlay media consisting of 2X MEM with 2X Penicillin/Streptomycin, 1mM HEPES, 2 µg/ml of TPCK and 3% Carboxy Methyl Cellulose (MP Biomedical) with 0.2 mg/mL DEAE-dextran (MP Biomedical) was added and well plates were incubated at 37 °C for 4 days. Plaques were fixed with 10% buffered formalin (Fisher), washed with DI water, and stained with 0.5% crystal violet (Thermo Fisher Scientific) for visualization.

## 3. Results

### 3.1 Single cell virus infection using drop-based microfluidics

A schematic of the virus infection workflow to compare drops to bulk is outlined in Fig 1. Naïve cells were infected with IAV at an MOI of 0.1 prior to encapsulation such that most cells were infected with a single infectious virus particle (Fig. 1A). Following inoculation, cells were dissociated from the tissue culture plates and processed for encapsulation into 100 µm drops and bulk samples were replated while the remaining cells were encapsulated as aqueous drops (Fig. 1B). Both encapsulated and bulk cells were processed similarly to ensure that trypsinization from the tissue culture plates and subsequent centrifugation and washing did not interfere with infection kinetics. A “0” h bulk and drop sample was obtained immediately following replating or encapsulation. The remaining bulk and drop samples were incubated for 24 h at 37°C to allow for a minimum of one round of virus replication (Fig. 1C and D). At 24 hpi, viral supernatant was collected from both the bulk and drop samples for analysis of virus replication and production via qRT-PCR and plaque assays, respectively. To sample viral supernatant from encapsulated infections, the drops were placed in the freezer and, upon thawing, treated with 20% PFO in HFE7500 to break the emulsions and release the cells and viral supernatant for collection (Fig. 1E). We compared cell viability, viral replication and infectious virus production from cells maintained in bulk culture and from cells encapsulated in drops (Fig. 1F).

**Figure 1.**
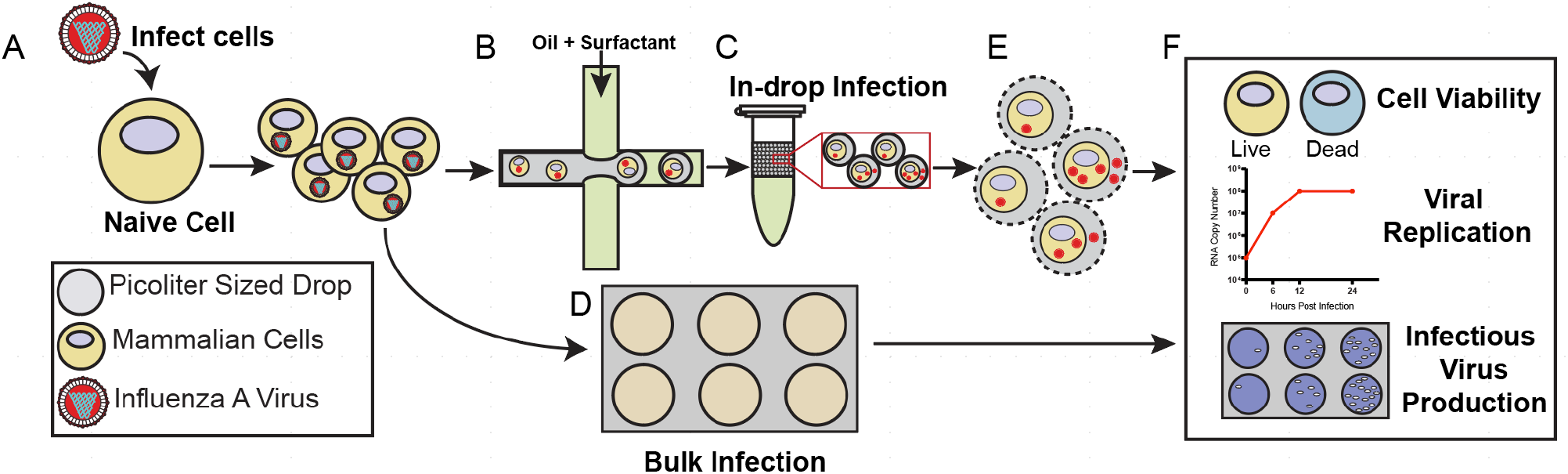
Graphical workflow. **(A)** Cells are infected with IAV **(B)** A suspension of cells is encapsulated into drops with a fluorinated continuous phase or replated onto standard tissue culture plates. **(C)** Drops are incubated for 24 h with standard tissue culture conditions. **(D)** A population of infected cells is replated onto standard tissue culture dishes, representing a bulk infection, and incubated for 24 h. **(E)** The emulsions are broken and cells and/or viral supernatant are recovered. **(F)** Cells and virus from in-drop and bulk infections are analyzed using live/dead staining to determine cell viability, RT-qPCR to determine viral copy number, and plaque assays to determine viral infectivity.

### 3.2 Validating IAV infection in different cell lines

Before performing IAV infection in drops, IAV viral replication was studied in bulk using three different cell lines: A549, MDCK, and Siat7e cells. Both the A549 and MDCK cells are anchorage dependent and are commonly used to propagate IAV and investigate various aspects of viral infection. Siat7e cells are a modified MDCK cells line that can grow in suspension and were developed to simplify cell culture-based IAV vaccine production (Chu et al., 2009; Chu et al., 2010). The Siat7e cells are anchorage independent and were tested for their potentially more favorable compatibility to drop encapsulation (Fischer et al., 2015; Tao et al., 2015a). IAV replication was compared between the different cell lines to determine the baseline kinetics of viral infection in a bulk infection. The cells were infected at an MOI of 0.1 and viral RNA abundance were measured at 0, 6, 12, 24, 30, 36 and 48 hpi (Fig. 2A-C). A549 cells demonstrated the most robust viral RNA replication during infection with 9.5 × 10^4^ genome copies/µL at 0 hpi, which increased to 3.9 × 10^8^ genome copies/µL at 24 hpi, a 1000-fold increase. Between 24 and 48 hpi, the amount of RNA detected fluctuated slightly but remained between 1.6 and 6.1 × 10^8^ genome copies/µL. MDCK cells demonstrated a slower increase in viral replication with 1.1 × 10^5^ genome copies/µL at 0 hpi, increasing to 3.5 × 10^7^ genome copies/µL at 24 hpi, a 100-fold increase, before reaching 1.9× 10^8^ genome copies/ µL at 48 hpi. IAV replication in Siat7e cells was limited with no exponential increase over the course of 48 hpi, with 6.0 × 10^5^ genome copies/µL at 0 hpi and 5.4 × 10^6^ genome copies/ µLat 48 hpi. The average genomes/µL for A549, MDCK, and Siat7e cells from 6-48 hpi, the time frame that represents active IAV replication where the 0 h measurement represents the inoculating dose,
was 2.6 × 10^8^, 6.4 × 10^7^, and 2.9 × 10^6^, respectively, demonstrating cell type dependent differences in IAV replication.

**Figure 2.**
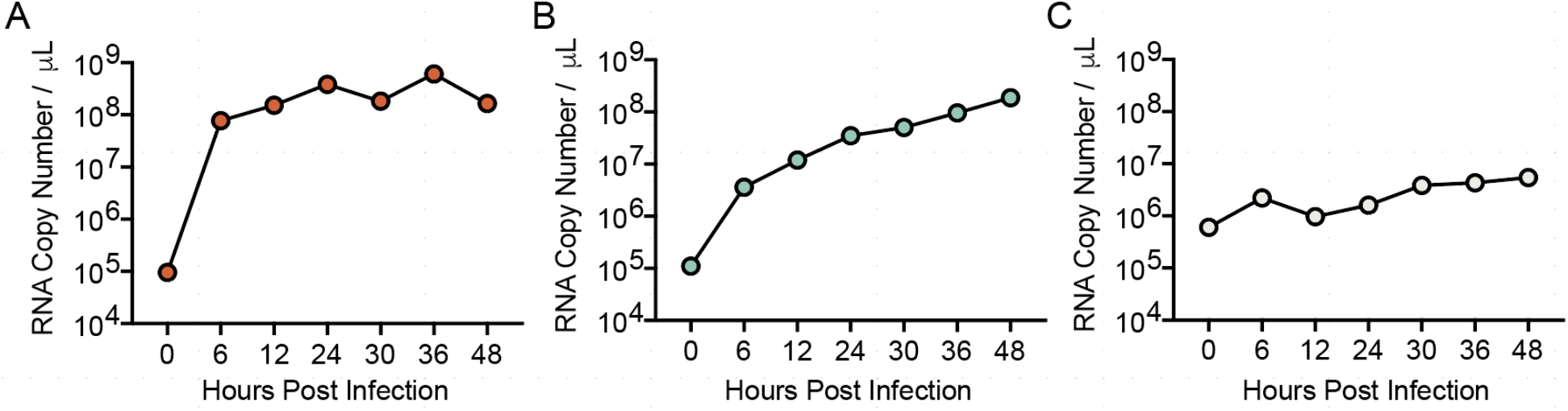
IAV infection in different cell lines. IAV replication, as measured by RT-qPCR over 48 hpi in **(A)** A549, **(B)** MDCK, and **(C)** Siat7e cells. All data represents the mean ± SD of a minimum of two independent replicates. Error bars are present.

### 3.3 Drop stability and cell viability during incubation

Studying viral infection of individual cells within microfluidic drops requires that drops resist coalescence following cell encapsulation and incubation. Cell encapsulation and incubation in different medias can change the aqueous chemistry of microfluidic drops, and increases in salt concentrations or temperature (37°C overnight) can disrupt drop stability (Etienne, Kessler, and Amstad, 2017). The A549, MDCK and Siat7e cells were encapsulated in 100 µm drops and incubated at 37 °C/5% CO2 incubator for 24 h. The distribution of drop radii (*R*) between 0 and 24 h was used to quantify drop stability following encapsulation and incubation of all three cell lines (Fig 3A). The average *R* at 0 h for A549 cells was 45.1 μm with a 95% confidence interval (CI) of [35.9, 54.3] which decreased by 0.9 μm with a 95% CI of [0.99, 0.75] μm after incubation for 24 h. For MDCK cells the average *R* at 0 h was 44.6 μm, with a 95% CI of [35.3, 53.9] μm, which increased by 1.32 μm with a 95% CI of [1.15, 1.49] μm after incubation for 24 h. For the Siat7e cells the average *R* at 0 h was 44.8 μm with a 95% CI of [35.5, 54.2] μm, and similar to MDCK cells this increased by 0.7 μm with a 95% CI of [0.57, 0.91] μm after incubation for 24 h. A linear mixed effects model, with the experimental date as a random factor, was used to determine if there was a significant change in drop *R* and if this change is a function of the cell type or timepoint post encapsulation. The change in drop *R* from 0 to 24 h following encapsulation of either A549 or MDCK cells was considered significantly different (p-value = < 0.001 for both) while the change in drop *R* from encapsulated Siat7e cells was not significant (p-value = 0.323). However, given the small changes in drop *R* compared to the drop size and the changes in drop size being less than the camera resolution of 1.6 μm/pixel, these data indicated that cell incubation does not have a meaningful impact on changes in drop stability, as measured by changes in drop *R*. Representative images of encapsulated A549 cells at 0 h (Fig 3B) and 24 h (Fig 3C) further demonstrates drop stability.

**Figure 3.**
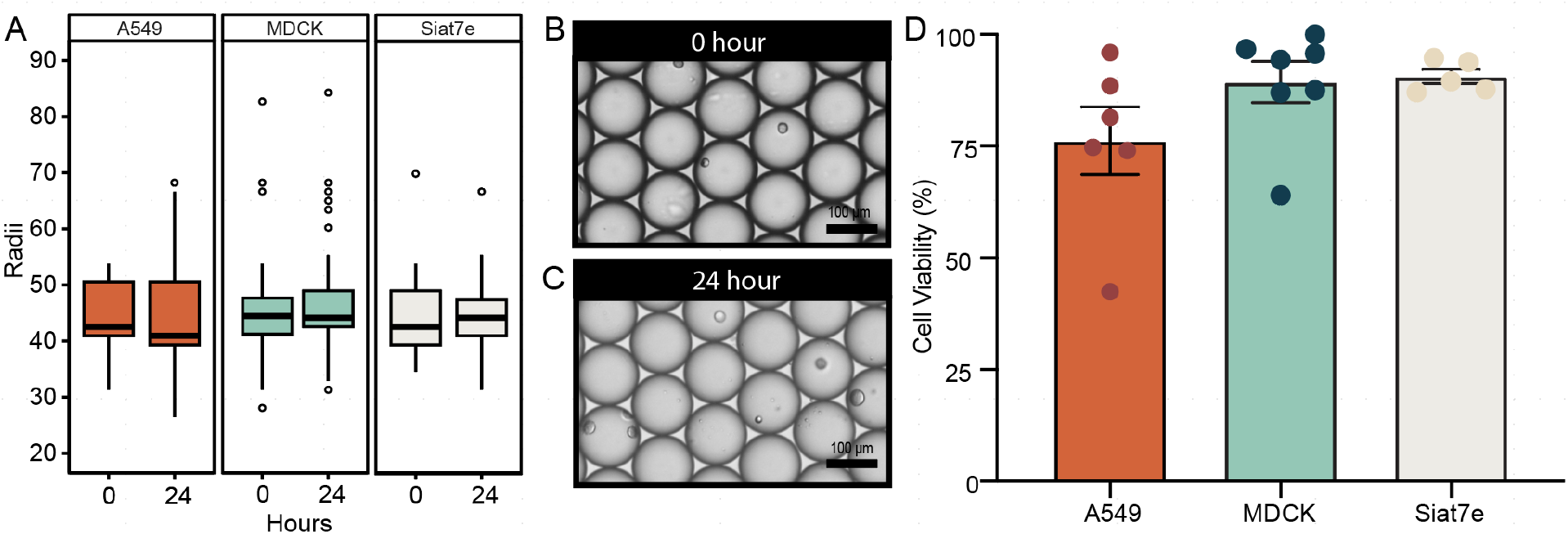
Drop stability and cell viability in 100 μm microfluidic drops. **(A)** Drop radii of encapsulated A549 (n=4,067), MDCK (n=4,235), and Siat7e (n=3,858) at 0 and 24 h. **(B)** Representative images of A549 cells encapsulated in drops at 0 h and **(C)** 24 h. Scale bars are 100 μm. **(D)** A549, MDCK, and Siat7e cell viability at 24 h post-encapsulation as represented by the mean ± SEM.

To assess cell viability, drops containing cells were broken after 24 hours of incubation and the aqueous layer which contained the cells was collected and analyzed with a colorimetric cell viability stain. Each of the three cell lines showed cell viability over 75%. Average live cell viability percentages were 76.2 ± 7.6% for A549 cells, 89.3 ± 4.6% for MDCK cells, and 90.6 ± 3.5% for Siat7e cells. The high cell viability observed in the Siat7e cells was expected as they are non-adherent so there are no physiological changes when encapsulated within drops. For the adherent cell lines, viability of the MDCK cells was similar to the Siat7e. While viability of the A549 cells was less than that observed in the MDCK and Siat7e cells in drops, there was not a significant difference. We hypothesize that the differences observed in viability between the adherent A549 and MDCK cells was due to inherent differences between the cells themselves. MDCK cells are more prone to overgrowing and detaching from tissue culture flasks and do not display strong contact inhibition and therefore may be slightly better suited for drop encapsulation. In comparison, A549 cells demonstrate strong contact inhibition and rarely detach from tissue culture flasks. These inherent differences may impact cell viability following encapsulation into microfluidic drops.

### 3.4 Cell loading in microfluidic drops

The majority of drops should contain one cell in single cell studies. The number of cells loaded into drops can be estimated using a Poisson distribution when the concentration of cells is known and the cells do not interact with each other during encapsulation (Collins et al., 2015). Thus, the average number of cells per drop (*λ*) can be estimated using the starting cell concentration and the drop volume (Mazutis et al., 2013). Loading cells at a density of 2 × 10^6^ cells/mL into 100 µm drops should result in a *λ* of 1.0 and result in 37% of drops containing no cells, 37% of drops containing one cell, 18% of drops containing two cells, 6% of drops containing three cells, and 1.9% of drops containing four or more cells (Poisson statistics). All three cell lines were therefore loaded at a density of 2 × 10^6^ cells/mL and analyzed to determine if they followed a *λ* of 1. Cell counts were completed using high-speed video footage at the microfluidic device flow-focusing junction as well as still images of drops loaded onto a hemocytometer (Fig 4A and 4B). A549 cells had 10.8 ± 1.9% and 13.1 ± 3.6% of drops containing one cell on the hemocytometer and high-speed camera, respectively (Fig 4C). MDCK cells had 10.2 ± 1.9% and 5.8 ± 2.0% of drops containing one cell on the hemocytometer and high-speed camera, respectively (Fig 4D). Siat7e cells had 5.7 ± 1.9% and 11.0 ± 1.3% of drops containing one cell on the hemocytometer and high-speed camera, respectively (Fig 4E).

**Figure 4:**
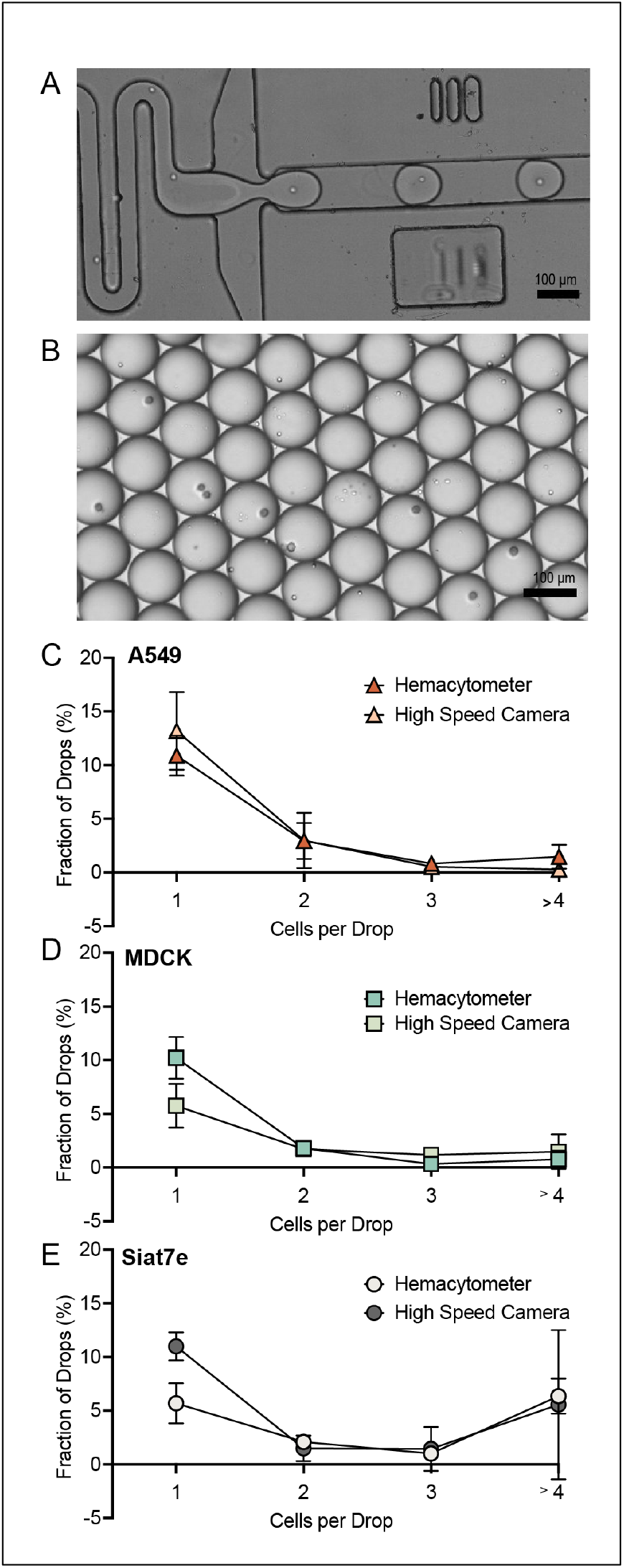
Cell encapsulation in microfluidic drops. **(A)** Still image of high-speed camera footage during cell encapsulation. **(B)** Representative image of drops encapsulated onto a hemocytometer. **(C)** A549 cell loading as measured with the hemocytometer (red triangle, n=600) and high-speed camera (peach triangle, n=405) **(D)** MDCK cell loading as measured with the hemocytometer (aqua square, n=601) and high-speed camera (green square, n=393) **(E)** Siat7e cell loading as measured with the hemocytometer (light grey circle, n=598) and high-speed camera (dark grey circle, n=431). All scale bars are 100 μm.

The calculated *λ* was used to compare the cell loading across the three cell lines and the two imaging methods to the theoretical Poisson distribution. The calculated *λ* ranged from 0.021 to 0.14, which corresponds to a theoretical Poisson distribution for a lower starting cell concentration of 3 × 10^5^ cells/mL. We hypothesize that this discrepancy was most likely due to cell settling in the syringe during loading and cell adherence to the filter upstream of the flow-focusing junction. However, the calculated *λ* depended on the measurement method and cell line being evaluated. For A549 cells, the *λ* was 0.021 with a 95% CI of [0.01, 0.043] when measured with the high-speed camera and was 0.041 with a 95% CI of [0.012, 0.126] when measured with a hemocytometer (p-value = 0.025). The *λ* for the Siat7e cells measured with the high-speed camera was 0.141 with a 95% CI of [0.044, 0.427] while the measurements on the same dates with a hemocytometer had a *λ* of 0.085 with a 95% CI of [0.027, 0.263] (p-value = 0.017). The drops containing the Siat7e cells had the widest 95% CI. These results show a significant difference in the measured cell loading for A549 cells and Siat7e cells as determined with the high-speed camera and the hemocytometer (p-values < 0.001). In contrast, the *λ* for MDCK cells was 0.019 with a 95% CI of [0.006, 0.066] when measured with the high-speed camera and was 0.025 with a 95% CI of [0.007, 0.085] when measured with a hemocytometer. This was not considered a significant difference (p-value = 0.942) suggesting that the two methods used for MDCK cells were comparable for determining cell loading.

Overall, the Siat7e cells demonstrated the greatest variability and unpredictability in cell loading across both methods. We hypothesize that this is due to the cells adhering together, which is also visible during normal cell growth in a shaker flask, in which large clumps of cells arise during growth. Due to poor cell encapsulation and low IAV replication during infection, Siat7e cells were excluded from further analysis.

### 3.5 IAV propagation in microfluidic drops

Following analysis of cell encapsulation and viability of A549 and MDCK cells, IAV propagation was compared between A549 and MDCK cells in bulk and in drops. A549 and MDCK cells were infected at an MOI of 0.1 with the A/California/07/2009 H1N1 IAV strain. Viral RNA, measured using RT-qPCR, and infectious virus output, determined by plaque assays, from cells in drops and cells incubated in bulk on standard tissue culture plates at 0 and 24 hpi allowed us to quantify the increase in virus replication and infectious virus output for both bulk and drop infections. For A549 cells, the number of genome copies/µL in drops, as measured by M gene RNA copy number using RT-qPCR, was 2.3 × 10^5^ genome copies/µL at 0 h and increased to 2.3 × 10^7^ genome copies/µL at 24 h (Fig 5A). Bulk infections of A549 cells had a comparable increase with 1.8 × 10^6^ genome copies at 0 h and 5.6 × 10^8^ genome copies/µL at 24 h (Fig 5A). The log RNA concentration in drops compared to bulk at 0 h and at 24 h were significantly different (p-value 0.0006 and 4.3E-09, respectively) which we hypothesize is due to a lower number of cells associated with the in-drop samples due to calculated cell encapsulation. However, the log difference of RNA produced by cells from 0 to 24 h in drops compared to cells in bulk was not significant (p-value 0.057) and suggests that viral replication was not impacted by cell encapsulation in drops. Recovery of infectious virus from A549 cells over the same 24 h incubation period was also consistent between IAV infection in drops and bulk culture with 1.1 × 10^5^ PFU/mL and 3.6 × 10^5^ PFU/mL recovered at 24 hpi, respectively (Fig 5B). Surprisingly, the log PFU/mL concentration in drops compared to bulk at 0 h and at 24 h was not significantly different (p-value 0.31 and 0.71, respectively). In addition, the amount of infectious virus produced from 0 to 24 h in both drop and bulk infections was also not significantly different (p-value 0.84). The genome to PFU ratio results in 1 PFU per 2.97 × 10^2^ genomes for A549 cells in drops and 1 PFU per 1.63 × 10^3^ genomes in bulk, suggesting that any significant difference in RNA concentration is not impacting the amount of virus being produced by A549 cells in drops.

**Figure 5:**
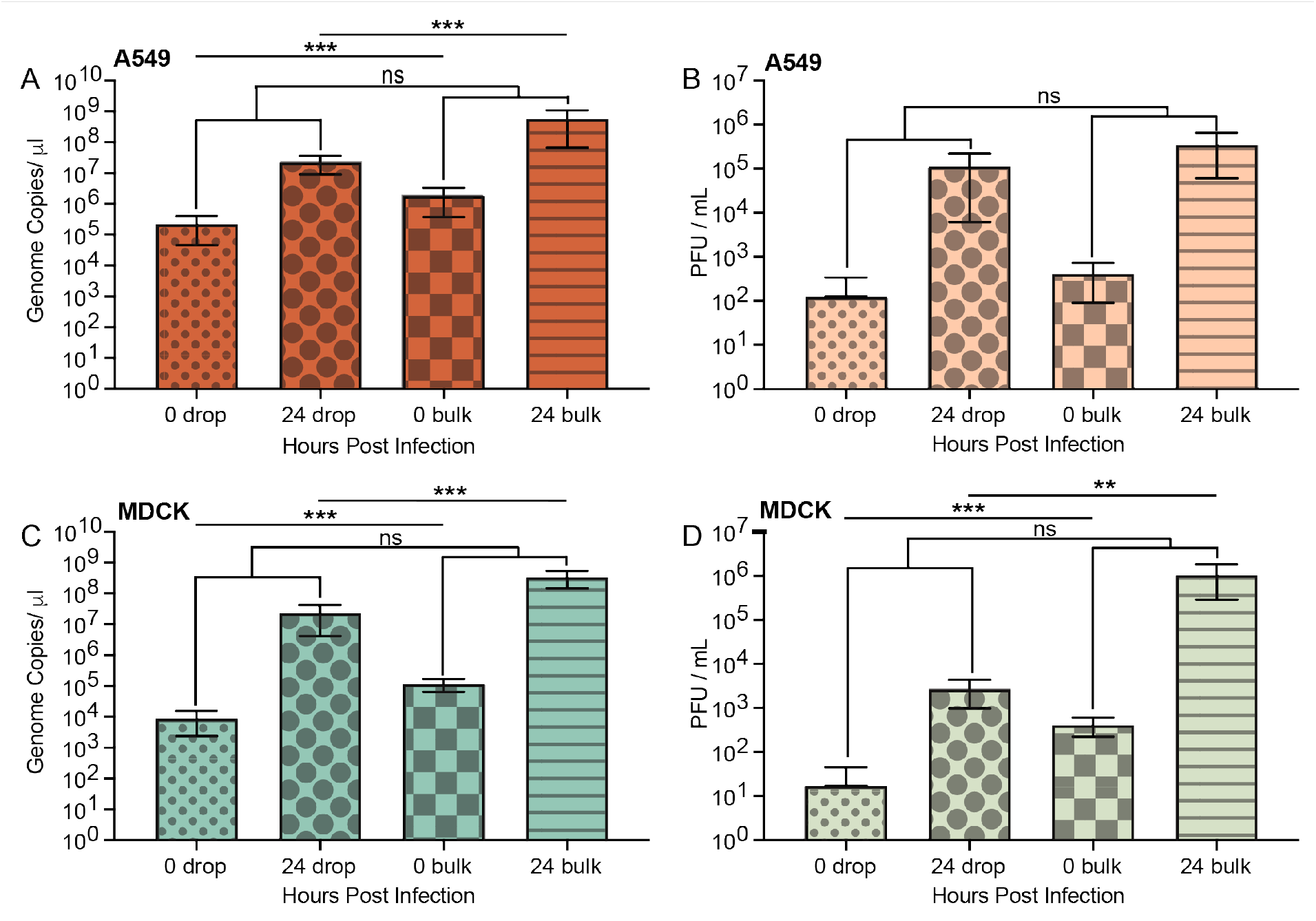
Comparison of in drop and bulk IAV infections in A549 and MDCK cells. **(A)** RNA copy number of A/Cal/07/2009 H1N1 IAV in A549 cells infected at an MOI of 0.1 at 0 hpi in drops (red small dots), 24 hpi in drops (red large dots), 0 hpi in bulk (red squares), and 24 hpi in bulk (red stripe). **(B)** Infectious virions (PFU/mL) released from A549 cells at 0 hpi in drops (peach small dots), 24 hpi in drops (peach large dots), 0 hpi in bulk (peach squares), and 24 hpi in bulk (peach stripe). **(C)** RNA copy number of A/Cal/07/2009 H1N1 IAV in MDCK cells infected at an MOI of 0.1 at 0 hpi in drops (aqua small dots), 24 hpi in drops (aqua large dots), 0 hpi in bulk (aqua squares), and 24 hpi in bulk (aqua stripe). **(D)** Infectious virions (PFU/mL) released from MDCK cells 0 hpi in drops (green small dots), 24 hpi in drops (green large dots), 0 hpi in bulk (green squares), and 24 hpi in bulk (green stripe). All data represented as the mean± the SD with a minimum of three independent experiments.

For MDCK cells, the number of genome copies in drops was 8.9 × 10^3^ genome copies/µL at 0 h and increased to 2.3 × 10^7^ genome copies/µL at 24 h. Whereas, bulk infections of MDCK cells increased from 1.2 × 10^5^ genome copies/µL at 0 h and 3.4 × 10^8^ genome copies/µL at 24 h (Fig 5C). Similar to A549 cells, the log RNA concentrations measured in MDCK cells in drops compared to bulk at 0 h and at 24 h were significantly different (p-value 7.2E-05 and 0.0002, respectively) while the overall log difference of RNA produced by cells from 0 to 24 h was not significantly different (p-value 0.6) between in drop and bulk infection. Recovery of infectious virus from MDCK cells over 24 h in drops was 2.6 × 10^3^ PFU/mL while bulk infections produced 1.0 × 10^6^ PFU/mL (Fig 5D). In contrast to A549 cells, the log PFU/mL concentrations measured in MDCK cells in drops compared to bulk at 0 h and at 24 h were significantly different (p-value 0.0004 and 0.003, respectively), however this did not translate into a significant difference in the change in log PFU/mL concentration from 0 to 24 h between in drop and bulk infections. The genome to PFU ratio for MDCK cells was 1 PFU per 5.81 × 10^3^ genomes in drops and 1 PFU per 3.24 × 10^2^ genomes in bulk. The low PFU to genome ratio in MDCK cells in drops could be the result of less virus being assembled compared to bulk culture whereas RNA replication appears to not be impacted by encapsulation.

## 4. Discussion

Microfluidics and single cell sequencing are expanding the field of single cell virology and enabling higher resolution analysis of the effects of cellular heterogeneity on viral infection dynamics. Recent studies in single cell virology include the kinetics of viral production (Akpinar, Timm, and Yin, 2016; Guo et al., 2017; Timm and Yin, 2012; Zhu, Yongky, and Yin, 2009), the distribution of viral burst size (Schulte and Andino, 2014), and characterization of genetic variability using single cell-RNA sequencing (Drayman et al., 2019; Russell et al., 2019; Russell et al., 2018). To understand how viral and/or cellular heterogeneity impacts the outcomes of viral infection at a single cell level requires the ability to interrogate large populations of individually infected cells. Many of the current studies depend upon diluting populations of cells into standard well-plates to assess single cells, isolation of individual cells by FACS, or isolating cells in microfluidic chambers (Guo et al., 2017). However, analyses in well-plates is typically is limited to hundreds of single cells, while 2D microfluidic devices are limited to a few thousand cells (Combe et al., 2015; Guo et al., 2017; Heldt et al., 2015; Kupke et al., 2020). To process tens of thousands or up to millions of single cells, a continuous flow technique such as FACS or drop-based microfluidics is necessary for much higher throughput, thousands of cells per second (Drayman et al., 2019; Sun et al., 2020). One drawback in isolating infected cells by FACS is it requires the use of antibodies to label infected cells or fluorescent markers encoded within the virus itself, which may not be accurate in highly heterogenous populations. FACS sorting can also be limited to early timepoints post infection to ensure that virus spread is reduced, and infection parameters are similar so that data is comparable from cell to cell. FACS analysis of infected cells at early time points limits the ability to investigate heterogeneity at later time points post infection and an inability to explore virus production. In comparison, drop-based microfluidics can be used to compartmentalize individual cells within drops, thereby creating up to millions of tiny bioreactors for further downstream analysis (Rotem et al., 2018; Tao et al., 2015a). These compartmentalized cells in drops allow for viral infection to proceed without virus spread to neighboring drops, providing a way to analyze how infection proceeds at a multitude of time points and the ability to analyze virus replication, production and cellular responses at a single cell level. To date, drop-based viral infections have only been performed with MNV-1 using cell lines adapted to spinner cultures or already grown in suspension (Fischer et al., 2015; Tao et al., 2015a; Tao et al., 2015b). The use of spinner or suspension-based cell lines for drop-based microfluidics limits the applicability of the technology for studying viral infections that predominantly occur in adherent cell culture-based models. In addition, studying viral infections within drops has not been extended to include the plethora of eukaryotic viruses that exist and impact our daily lives. The ability to expand drop-based microfluidics to include a range of different cell lines and viruses would allow for more in-depth analysis of viral infection with single cell resolution. Our approach evaluated the ability of standard cell lines to propagate IAV infection at a single cell level using drop-based microfluidics.

We found that 100 μm drops loaded with the A549, MDCK or Siat7e cells remained stable following overnight incubation and cell viability remained high. High cell viability was expected from the Siat7e cells as they are grown in shaker flasks in suspension. The high cell viability from the A549 and MDCK cells, while more variable between experiments than the Siat7e cells, was also promising as both cell lines are adherent cell lines traditionally grown on tissue culture flasks and vessels. Previous studies exploring viability of cells in drops found that smaller drop sizes resulted in low levels of cell viability, even after a couple of hours, most likely due to lack of enough nutrients or buildup of waste products within the drop (Köster et al., 2008). Our data demonstrates that drop-based microfluidics can be used to encapsulate and incubate a wider range of cell types for single cell analysis. While we only analyzed cells over a 24h time span to accommodate the lifecycle of IAV, we expect that incubation of encapsulated cells for longer periods is possible.

The majority of drops with cells contained 1 cell per drop after loading. Using a Poisson distribution to model the expected cell loading based on cell concentration and drop size, we determined that a *λ* of 1, in which the majority of drops contained 1 cell, was represented by a cell concentration of 2 × 10^6^ cells/mL for 100 μm drops (50 μm radii). We also assumed that all the cells in the suspension have an equal probability of encapsulation. However, throughout our experiments, cells settled in the syringes and attached to the aqueous-stream filter on the microfluidic drop-maker, lowering the number of cells being encapsulated. Therefore, the *λ* in our experiments was similar to a much lower cell concentration, although the majority of our loaded drops still contained a single cell. Surprisingly, the *λ* for the Siat7e cell line had the largest variation and was significantly different from the other two cell lines, with more drops having with 4+ cells/drop than was observed for the A549 or MDCK cells. We hypothesize that this was due to the Siat7e cells adhering to each other when cultured in a shaker flask and during infection prior to cell encapsulation. To improve cell loading, a higher concentration of cells can be used, although previous analysis of 2, 4 or 8 × 10^6^ cells/mL in 50 μm drops had similar loading patterns (Fischer et al., 2015). We have observed cell loading that more closely follows the Poisson distribution for a given cell concentration with different cell types, such as true suspension cell lines (data not shown), so it is advisable to test cell loading for any new cell types being encapsulated into microfluidic drops. Furthermore, our analysis methods using either a high-speed camera or low-tech hemocytometer to determine cell loading resulted in significantly different *λ* values, with the low-tech hemocytometer providing more consistent results. This suggests that while either method can be utilized for quantifying cell loading, a simple count using images taken on a hemocytometer is sufficient.

Following our success in encapsulation and culturing multiple cell lines within microfluidic drops, we investigated whether infected cells would continue to produce virus following encapsulation. We explored the ability of A549 and MDCK cells only due to highly variable cell loading and low virus production from the Siat7e cells. Viral RNA and titers were measured at 0 and 24 h post encapsulation and compared to standard bulk infections on well plates. The amount of virus produced from each cell line between 0 and 24 h in bulk was not significantly different than the amount of virus produced from cells encapsulated and incubated within drops. Our data demonstrates that microfluidic drops can be used to propagate IAV from individual cells and that adherent standard cell lines can be used, reducing the need for development of different model systems for studying infections at a single cell level.

Previous analysis of virus infection in drops was performed with MNV-1, a positive sense RNA virus in a small icosahedral capsid. In comparison to IAV, MNV-1 is extremely stable at a range of environmental conditions and lacks an outer envelope (Henderson, 2008). Here we have demonstrated that an enveloped virus with reduced environmental stability can be produced from standard cell lines in drops at a similar rate to what is observed in a bulk culture, which enables more extensive analysis of virus infection at a single cell level. Previous studies exploring viral kinetics and replication (Holmes, Zhang, and Bieniasz, 2015; Timm and Yin, 2012) and innate immune activation (Russell et al., 2019; Timm et al., 2017), have provided new insight into the heterogeneity surrounding viral infections. The ability to expand single cell analysis to tens of thousands of cells in a high throughput manner has the potential to revolutionize virology studies.

## 5. Conclusions

Analysis of individually infected cells has revealed how complex and heterogenous viral infections are, even in controlled laboratory settings (Combe et al., 2015; Russell et al., 2019; Russell et al., 2018; Sun et al., 2020). The implementation of -omics technology to these systems has further demonstrated how viral infections are as complicated at a single cell level as they are within a larger host or system (Prakadan et al., 2017; Xu et al., 2020; Zilionis et al., 2017). These applications of new technology to the study of virology allow us to look closer at how individual host cells drive viral evolution and responses to the innate and adaptive immune systems. With the current COVID-19 pandemic and the emergence of multiple viral variants with populations, it is critical to expand our ability to study how these variants arise, how they operate under selective pressures, and how this impacts transmission and virulence moving forward (Harvey et al., 2021). The ability to study these aspects in individual cells has been limited by low throughput analysis, reliance on expensive cell sorting machinery, or viruses that are easily manipulated to express fluorescent markers. The use of drop-based microfluidics offers the ability to perform high throughput analysis of individual cells (Gérard et al., 2020; Loveday et al., 2021; Matuła et al., 2020). In addition, these applications can be adapted to many laboratories with minimal investment. Our approach and methods described here provide a framework for pursuing single cell studies of virus infections using drop-based microfluidics.

## References

Akpinar, F., Timm, A. and Yin, J., 2016. High-throughput single-cell kinetics of virus infections in the presence of defective interfering particles. Journal of virology 90, 1599–1612.

Anna, S.L., Bontoux, N. and Stone, H.A., 2003. Formation of dispersions using “flow focusing” in microchannels. Applied Physics Letters 82, 364–366.

Brooke, C.B., 2017. Population Diversity and Collective Interactions during Influenza Virus Infection. J Virol 91.

Chu, C., Lugovtsev, V., Golding, H., Betenbaugh, M. and Shiloach, J., 2009. Conversion of MDCK cell line to suspension culture by transfecting with human siat7e gene and its application for influenza virus production. Proc. Natl. Acad. Sci. U. S. A. 106, 14802–14807.

Chu, C., Lugovtsev, V., Lewis, A., Betenbaugh, M. and Shiloach, J., 2010. Production and antigenic properties of influenza virus from suspension MDCK-siat7e cells in a bench-scale bioreactor. Vaccine 28, 7193–7201.

Collins, D.J., Neild, A., deMello, A., Liu, A.Q. and Ai, Y., 2015. The Poisson distribution and beyond: methods for microfluidic droplet production and single cell encapsulation. Lab Chip 15, 3439–59.

Combe, M., Garijo, R., Geller, R., Cuevas, J.M. and Sanjuán, R., 2015. Single-Cell Analysis of RNA Virus Infection Identifies Multiple Genetically Diverse Viral Genomes within Single Infectious Units. Cell Host Microbe 18, 424–432.

Cristinelli, S. and Ciuffi, A., 2018. The use of single-cell RNA-Seq to understand virus-host interactions. Curr. Opin. Virol. 29, 39–50.

Dolan, P.T., Whitfield, Z.J. and Andino, R., 2018. Mapping the Evolutionary Potential of RNA Viruses. Cell Host Microbe 23, 435–446.

Drayman, N., Patel, P., Vistain, L. and Tay, S., 2019. HSV-1 single-cell analysis reveals the activation of anti-viral and developmental programs in distinct sub-populations. Elife 8.

Duffy, D.C., McDonald, J.C., Schueller, O.J.A. and Whitesides, G.M., 1998. Rapid Prototyping of Microfluidic Systems in Poly(dimethylsiloxane). Analytical Chemistry 70, 4974–4984.

Etienne, G., Kessler, M. and Amstad, E., 2017. Influence of Fluorinated Surfactant Composition on the Stability of Emulsion Drops. Macromolecular Chemistry and Physics 218.

Fischer, A.E., Wu, S.K., Proescher, J.B.G., Rotem, A., Chang, C.B., Zhang, H., Tao, Y., Mehoke, T.S., Thielen, P.M., Kolawole, A.O., Smith, T.J., Wobus, C.E., Weitz, D.A., Lin, J.S., Feldman, A.B. and Wolfe, J.T., 2015. A high-throughput drop microfluidic system for virus culture and analysis. J. Virol. Methods 213, 111–117.

Gérard, A., Woolfe, A., Mottet, G., Reichen, M., Castrillon, C., Menrath, V., Ellouze, S., Poitou, A., Doineau, R., Briseno-Roa, L., Canales-Herrerias, P., Mary, P., Rose, G., Ortega, C., Delincé, M., Essono, S., Jia, B., Iannascoli, B., Richard-Le Goff, O., Kumar, R., Stewart, S.N., Pousse, Y., Shen, B., Grosselin, K., Saudemont, B., Sautel-Caillé, A., Godina, A., McNamara, S., Eyer, K., Millot, G.A., Baudry, J., England, P., Nizak, C., Jensen, A., Griffiths, A.D., Bruhns, P. and Brenan, C., 2020. High-throughput single-cell activity-based screening and sequencing of antibodies using droplet microfluidics. Nat. Biotechnol.

Guo, F., Li, S., Caglar, M.U., Mao, Z., Liu, W., Woodman, A., Arnold, J.J., Wilke, C.O., Huang, T.J. and Cameron, C.E., 2017. Single-Cell Virology: On-Chip Investigation of Viral Infection Dynamics. Cell Rep. 21, 1692–1704.

Harvey, W.T., Carabelli, A.M., Jackson, B., Gupta, R.K., Thomson, E.C., Harrison, E.M., Ludden, C., Reeve, R., Rambaut, A., Consortium, C.-G.U., Peacock, S.J. and Robertson, D.L., 2021. SARS-CoV-2 variants, spike mutations and immune escape. Nat Rev Microbiol 19, 409–424.

Heldt, F.S., Kupke, S.Y., Dorl, S., Reichl, U. and Frensing, T., 2015. Single-cell analysis and stochastic modelling unveil large cell-to-cell variability in influenza A virus infection. Nat. Commun. 6, 8938.

Henderson, K.S., 2008. Murine norovirus, a recently discovered and highly prevalent viral agent of mice. Lab Anim (NY) 37, 314–20.

Holmes, M., Zhang, F. and Bieniasz, P.D., 2015. Single-Cell and Single-Cycle Analysis of HIV-1 Replication. PLoS Pathog. 11, e1004961.

Holtze, C., Rowat, A.C., Agresti, J.J., Hutchison, J.B., Angile, F.E., Schmitz, C.H., Koster, S., Duan, H., Humphry, K.J., Scanga, R.A., Johnson, J.S., Pisignano, D. and Weitz, D.A., 2008. Biocompatible surfactants for water-in-fluorocarbon emulsions. Lab Chip 8, 1632–9.

Köster, S., Angilè, F.E., Duan, H., Agresti, J.J., Wintner, A., Schmitz, C., Rowat, A.C., Merten, C.A., Pisignano, D., Griffiths, A.D. and Weitz, D.A., 2008. Drop-based microfluidic devices for encapsulation of single cells. Lab Chip 8, 1110–1115.

Kupke, S.Y., Ly, L.-H., Börno, S.T., Ruff, A., Timmermann, B., Vingron, M., Haas, S. and Reichl, U., 2020. Single-Cell Analysis Uncovers a Vast Diversity in Intracellular Viral Defective Interfering RNA Content Affecting the Large Cell-to-Cell Heterogeneity in Influenza A Virus Replication. Viruses 12.

Liao, M., Liu, Y., Yuan, J., Wen, Y., Xu, G., Zhao, J., Cheng, L., Li, J., Wang, X., Wang, F., Liu, L., Amit, I., Zhang, S. and Zhang, Z., 2020. Single-cell landscape of bronchoalveolar immune cells in patients with COVID-19. Nat. Med. 26, 842–844.

Lin, J., Jordi, C., Son, M., Van Phan, H., Drayman, N., Abasiyanik, M.F., Vistain, L., Tu, H.-L. and Tay, S., 2019. Ultra-sensitive digital quantification of proteins and mRNA in single cells. Nat. Commun. 10, 3544.

Loveday, E.K., Zath, G.K., Bikos, D.A., Jay, Z.J. and Chang, C.B., 2021. Screening of Additive Formulations Enables Off-Chip Drop Reverse Transcription Quantitative Polymerase Chain Reaction of Single Influenza A Virus Genomes. Anal Chem 93, 4365–4373.

Matuła, K., Rivello, F. and Huck, W.T.S., 2020. Single-Cell Analysis Using Droplet Microfluidics. Adv Biosyst 4, e1900188.

Mazutis, L., Gilbert, J., Ung, W.L., Weitz, D.A., Griffiths, A.D. and Heyman, J.A., 2013. Single-cell analysis and sorting using droplet-based microfluidics. Nat. Protoc. 8, 870–891.

Petrova, V.N. and Russell, C.A., 2018. The evolution of seasonal influenza viruses. Nat. Rev. Microbiol. 16, 47–60.

Prakadan, S.M., Shalek, A.K. and Weitz, D.A., 2017. Scaling by shrinking: empowering single-cell ‘omics’ with microfluidic devices. Nat. Rev. Genet. 18, 345–361.

Rotem, A., Serohijos, A.W., Chang, C.B., Wolfe, J.T., Fischer, A.E., Mehoke, T.S., Zhang, H., Tao, Y., Lloyd Ung, W. and Choi, J.-M., 2018. Evolution on the biophysical fitness landscape of an RNA virus. Molecular biology and evolution 35, 2390–2400.

Russell, A.B., Elshina, E., Kowalsky, J.R., te Velthuis, A.J. and Bloom, J.D., 2019. Single-cell virus sequencing of influenza infections that trigger innate immunity. Journal of virology, JVI. 00500–19.

Russell, A.B., Trapnell, C. and Bloom, J.D., 2018. Extreme heterogeneity of influenza virus infection in single cells. Elife 7.

Schulte, M.B. and Andino, R., 2014. Single-cell analysis uncovers extensive biological noise in poliovirus replication. Journal of virology 88, 6205–6212.

Sen, A., Rothenberg, M.E., Mukherjee, G., Feng, N., Kalisky, T., Nair, N., Johnstone, I.M., Clarke, M.F. and Greenberg, H.B., 2012. Innate immune response to homologous rotavirus infection in the small intestinal villous epithelium at single-cell resolution. Proc. Natl. Acad. Sci. U. S. A. 109, 20667–20672.

Shu, B., Wu, K.H., Emery, S., Villanueva, J., Johnson, R., Guthrie, E., Berman, L., Warnes, C., Barnes, N., Klimov, A. and Lindstrom, S., 2011. Design and performance of the CDC real-time reverse transcriptase PCR swine flu panel for detection of 2009 A (H1N1) pandemic influenza virus. J Clin Microbiol 49, 2614–9.

Sun, J., Vera, J.C., Drnevich, J., Lin, Y.T., Ke, R. and Brooke, C.B., 2020. Single cell heterogeneity in influenza A virus gene expression shapes the innate antiviral response to infection. PLoS Pathog 16, e1008671.

Tao, Y., Rotem, A., Zhang, H., Chang, C.B., Basu, A., Kolawole, A.O., koehler, S., Ren, Y., Lin, J.S., Pipas, J.M., Feldman, A.B., Wobus, C. and Weitz, D.A., 2015a. Rapid, targeted and culture-free viral infectivity assay in drop-based microfluidics. Lab on a Chip.

Tao, Y., Rotem, A., Zhang, H., Cockrell, S.K., Koehler, S.A., Chang, C.B., Ung, L.W., Cantalupo, P.G., Ren, Y., Lin, J.S., Feldman, A.B., Wobus, C.E., Pipas, J.M. and Weitz, D.A., 2015b. Artifact-Free Quantification and Sequencing of Rare Recombinant Viruses by Using Drop-Based Microfluidics. Chembiochem 16, 2167–2171.

Timm, A. and Yin, J., 2012. Kinetics of virus production from single cells. Virology 424, 11–17.

Timm, A.C., Warrick, J.W. and Yin, J., 2017. Quantitative profiling of innate immune activation by viral infection in single cells. Integr. Biol. 9, 782–791.

Weingartl, H.M., Berhane, Y., Hisanaga, T., Neufeld, J., Kehler, H., Emburry-Hyatt, C., Hooper-McGreevy, K., Kasloff, S., Dalman, B., Bystrom, J., Alexandersen, S., Li, Y. and Pasick, J., 2010. Genetic and pathobiologic characterization of pandemic H1N1 2009 influenza viruses from a naturally infected swine herd. Journal of virology 84, 2245–56.

Xu, X., Wang, J., Wu, L., Guo, J., Song, Y., Tian, T., Wang, W., Zhu, Z. and Yang, C., 2020. Microfluidic Single-Cell Omics Analysis. Small 16, e1903905.

Zanini, F., Pu, S.-Y., Bekerman, E., Einav, S. and Quake, S.R., 2018. Single-cell transcriptional dynamics of flavivirus infection. Elife 7.

Zhu, Y., Yongky, A. and Yin, J., 2009. Growth of an RNA virus in single cells reveals a broad fitness distribution. Virology 385, 39–46.

Zilionis, R., Nainys, J., Veres, A., Savova, V., Zemmour, D., Klein, A.M. and Mazutis, L., 2017. Single-cell barcoding and sequencing using droplet microfluidics. Nat. Protoc. 12, 44–73.

